# Effects of Curative-Intent Lung Cancer Therapy on Functional Exercise Capacity and Patient-Reported Outcomes

**DOI:** 10.1101/508234

**Authors:** Duc Ha, Andrew L. Ries, Scott M. Lippman, Mark M. Fuster

**Author notes:** Correspondence: Duc Ha, MD, MAS (ORCID 0000-0002-3003-9438); Institute for Health Research; Kaiser Permanente Colorado; 2550 S. Parker Rd Suite 200; Aurora, CO 80014, USA.

## Abstract

**Background:** Lung cancer treatment can lead to negative health consequences. We analyzed the effects of curative-intent lung cancer treatment on functional exercise capacity (EC) and patient-reported outcomes (PROs).

**Methods:** We performed a prospective-observational cohort study of consecutive patients with stage I-IIIA lung cancer undergoing curative-intent therapy and assessed functional EC [*primary* outcome, six-minute walk distance (6MWD)], cancer-specific quality of life (QoL) [*secondary* outcome, European Organization for Research and Treatment of Cancer QoL Questionnaire Core 30 (EORTC-QLQ-C30) summary score], and *exploratory* outcomes including dyspnea [University of California San Diego Shortness of Breath Questionnaire (UCSD-SOBQ)] and fatigue [Brief Fatigue Inventory (BFI)] symptoms before and at 1 to 3 months post-treatment. We analyzed the time effect of treatment on outcomes using multivariable generalized estimating equations.

**Results:** In 35 enrolled participants, treatment was borderline-significantly associated with a clinically-meaningful decline in functional EC [(mean change, 95% CI) 6MWD = −25.4m (−55.3, +4.47), p=0.10], non-significant change in cancer-specific QoL [EORTC-QLQ-C30 summary score = − 3.4 (−9.8, +3.0), p=0.30], and statistically-significant and clinically-meaningful higher dyspnea [UCSD-SOBQ = +13.1 (+5.7, +20.6), p=0.001] and increased fatigue [BFI = +10.0 (+2.9, +17.0), p=0.006].

**Conclusions:** Among the first prospective analysis of the effect of curative-intent lung cancer treatment on functional EC and PROs, we observed worsening dyspnea and fatigue, and possibly a decline in functional EC but not cancer-specific QoL at 1 to 3 months post-treatment. Interventions to reduce treatment-related morbidities and improve lung cancer survivorship may need to focus on reducing dyspnea, fatigue, and/or improving functional EC.

**Consent and Approval:** Written informed consent was obtained from each participant included in this study. All human investigations were performed after approval by the VA San Diego Healthcare System institutional review board and in accord with an assurance filed with and approved by the U.S. Department of Health and Human Services.

## Introduction

Up to 50% of patients with non-small cell lung cancer (NSCLC) are diagnosed with stage I-IIIA disease [1] and eligible to undergo curative-intent therapy through a combination of lung cancer resection surgery [2], definitive radiation [3], or concurrent chemoradiation [4]. Immediately following curative-intent therapy, lung cancer patients are at risk for worsening health due to the toxicities and side effects of treatment. Depending on the extent of resection, a loss of 10-15% of lung function [including forced expiratory volume in 1 second (FEV1) and diffusion capacity of the lung for carbon monoxide (DLCO)] is expected at 3-6 months [5, 6] following lung cancer resection surgery and may persist at one year [7, 8]. In addition, perioperative pulmonary [9] and cardiopulmonary [10] complications occur in 15% and 35% of patients, respectively, and can lead to negative health consequences beyond the perioperative period [11]. In those undergoing definitive radiation, 5-15% will develop radiation pneumonitis [12] and associated symptoms (e.g. cough, dyspnea, pleuritic chest pain). Patients undergoing chemotherapy including adjuvant therapy can experience neutropenia, cardiac ischemia, heart failure (HF), and worsening symptoms related to gastrointestinal side effects, fatigue, and neuropathy [13, 14]. Also, lung cancer patients have major comorbidities including chronic obstructive pulmonary disease (COPD, present in approximately 50% of patients) due to tobacco exposure and heart failure (HF, approximately 13%) [15], the health effects of which can be exacerbated by lung cancer treatment.

The identification and quantification of peri-treatment changes in health may identify important decrements which can be prevented and/or minimized to improve lung cancer morbidity and mortality. In addition, peri-treatment efforts to optimize cardiopulmonary function and reduce symptom burden may improve lung cancer survivors’ quality of life (QoL) and survival. In this project, we assessed the changes in health as reflected by functional exercise testing and patient-reported outcomes (PROs). We hypothesized that curative-intent therapy of stage I-IIIA lung cancer is associated with decrements in health as reflected by functional exercise capacity (EC) and cancer-specific QoL.

## Methods

### Study Overview

We performed a prospective, observational cohort study of patients undergoing curative-intent therapy for lung cancer. We identified eligible patients from a weekly list of consecutive cases presented at the VA San Diego Healthcare System (VASDHS) chest tumor board (CTB). We included adult lung cancer patients with clinical stage I-IIIA disease who are recommended by the CTB to undergo lung cancer resection surgery, definitive radio-ablation, or concurrent chemoradiation as the primary mode of treatment. We excluded patients undergoing concurrent systemic therapy for other cancers or those physically unable to perform functional EC evaluation (e.g. quadriplegia or amputees) (Figure 1).

**Figure 1:**
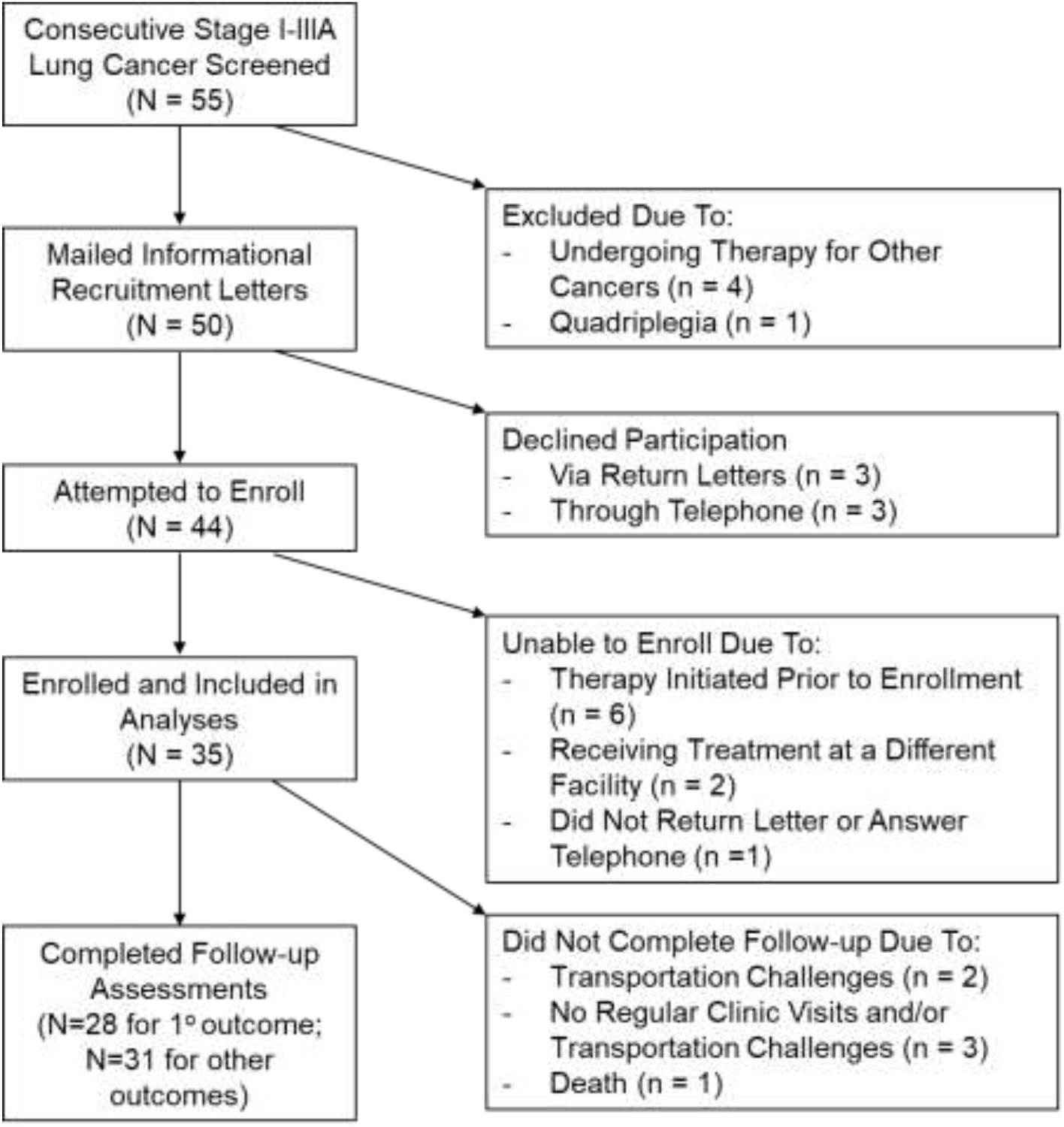
Flow Diagram of Enrolled Participants.

Between August 2016 and March 2018, we mailed informational letters to eligible patients after CTB management plans were communicated to patients and followed up with a telephone call approximately one week later to gauge their interest. Patients who were interested and willing to participate in our study were scheduled in-person visits, during which all provided written informed consent prior to study procedures. Outcome assessments were performed between August 2016 and May 2018, both before and at 1 to 3 months after completion of therapy. We followed the *STROBE* guideline recommendations to report our findings [16] and described our collection of potential confounders in the online data supplements. The VASDHS Institutional Review Board approved this protocol (#H150158).

### Functional EC and PRO Assessments

Our *primary* endpoint was functional EC as reflected by the six-minute walk test (6MWT) distance (6MWD). We chose the 6MWT [17] based on practical considerations of availability, ease of performance for testing, and the likelihood that daily activities of living are performed at submaximal exercise intensity. Also, the 6MWD is a valid measure of cardiorespiratory fitness in cancer [18] including lung cancer [19] clinical populations and has a defined minimal clinically important difference (MCID) cutoff available for interpretation (22 – 42 m, or a change of 9.5%) [20]. We performed the 6MWT according to the standard protocol at the VASDHS which follows the American Thoracic Society (ATS) Pulmonary Function Standards Committee recommendations [21].

Our *secondary* endpoint was a validated composite score of cancer-specific QoL, assessed by the European Organization for Research and Treatment of Cancer QoL Questionnaire Core 30 (EORTC-QLQ-C30) summary score [22]. We chose the EORTC-QLQ-C30 [23] based on availability, inclusion of core domains of QoL and other subdomains relevant to lung cancer (e.g. dyspnea, fatigue, pain), and a validated summary score [22] for analysis. We also performed *exploratory* PRO assessments for lung cancer-specific symptoms, generic health, sleep quality, dyspnea, fatigue, and anxiety/depression using the EORTC-QLQ-Lung Cancer Module 13 (LC13) [24], EuroQoL-5 Dimensions/visual analogue scale (EQ-5D/VAS) [25], Pittsburgh Sleep Quality Index (PSQI) [26], University California San Diego Shortness of Breath Questionnaire (UCSD SOBQ) [27], Brief Fatigue Inventory (BFI) [28], and Hospital Anxiety and Depression Scale (HADS) [29] questionnaires, respectively. We administered all questionnaires in-person when possible and on printed forms without modifications; all questionnaires were scored as per their respective instruction manuals.

### Sample Size

We calculated sample sizes assuming a significance level of p <0.05 in two-tailed tests and 80% power to detect a difference in outcomes. For our primary endpoint, we calculated that a sample size of 29 participants will be needed, using a MCID in the 6MWD of 40 m and standard deviation (SD) of 74 m as reported by previous literature [20]. For our secondary endpoint, we calculated a sample size of 30 participants based on a decline of 9 EORTC-QLQ-C30 summary score points following surgical lung cancer treatment [30] and SD of 17 as reported by previous literature [22]. These calculations were performed using PS Power and Sample Size Calculations software, Version 3.0.

### Statistical Analyses

We summarized descriptive statistics as means and standard deviations for all continuous variables and as counts and percentages for categorical variables. All outcomes were recorded and analyzed as continuous variables. The 6MWD was interpreted using the reference equations in healthy adults [31] and PROs using reference values where available or cutoffs from existing literature [32]. We used the paired-sample t-tests and multivariable generalized estimating equations (GEE) models to assess and analyze the effects of time and/or treatment on outcomes. We chose GEE models based on recommendations that they generally provide better model fits compared to linear mixed effects models for studies with a relatively large sample size (N >30) and few follow-up assessments [33]. To identify potential confounders, we used univariable (UVA) and multivariable (MVA) linear regression analyses to assess the relationship between baseline characteristics and the outcome of interest. We performed MVAs using stepwise backward selection modeling including all baseline characteristics with p <0.20; those in the final MVA were selected to enter multivariable GEE models. We further selected for covariates included in the final GEE models using stepwise backwards selection and p-value cutoff <0.20. To investigate the effect of treatment on outcomes, the effect of time (pre-/post-treatment) was forced into the model regardless of statistical significance. We also performed a pre-specified subgroup analysis of stage I lung cancer patients to compare the effects of surgery vs definitive radio-ablation on outcomes.

All data were entered and managed using REDCap electronic data capture tools hosted at the UCSD Clinical and Translational Research Institute [34] and analyzed using IBM^®^ SPSS^®^ Statistics software version 24.0. We used beta coefficients (β) and 95% confidence intervals (CIs) to describe effect size and defined statistical significance as p <0.05 in two-tailed tests.

## Results

### Participants

We screened 55 stage I-IIIA lung cancer patients, mailed recruitment letters to 50 eligible, and had a final enrollment of 35 participants (Figure 1); their baseline characteristics are summarized in Table 1. Most had a tobacco exposure history (32 participants, 91%), COPD (25, 71%), and stage I disease (29, 83%). There were no significant differences in baseline clinical characteristics for participants compared to non-participants except for a higher proportion of non-participants having stage II-IIIA disease (E-Table 1).

**Table 1:** Participant Characteristics.

| Participant Characteristics<br>(VASDHS, 2016-2018) | Value<br>(N=35) |
| --- | --- |
| Age, years, mean (SD) | 68.6 (7.3) |
| Race, n (%) |  |
| Asian | 2 (6) |
| Black | 4 (11) |
| Hispanic | 2 (6) |
| White | 27 (77) |
| Male sex, n (%) | 34 (97) |
| BMI, kg/m<sup>2</sup>, mean (SD) | 25.9 (4.8) |
| Smoking history, n (%) |  |
| Current | 15 (43) |
| Former | 17 (49) |
| Never | 3 (9) |
| Pack years, mean (SD) | 53.3 (34.1) |
| Comorbidities, n (%) |  |
| Hypertension | 25 (71) |
| Hyperlipidemia | 21 (60) |
| Diabetes | 9 (26) |
| CKD | 5 (14) |
| Atrial arrhythmia | 8 (23) |
| CAD | 11 (31) |
| HFrEF<sup>†</sup> | 5 (14) |
| PVD | 6 (17) |
| COPD | 25 (71) |
| OSA | 4 (11) |
| Anxiety/depression/PTSD | 9 (26) |
| Other cancer | 10 (29) |
| Lung function, mean (SD) |  |
| FEV<sub>1</sub>/FVC % | 59.8 (14.7) |
| FEV<sub>1</sub> % predicted | 70.0 (24.1) |
| TLC % predicted<sup>*</sup> | 113.3 (19.0) |
| DL<sub>CO</sub> % predicted | 80.4 (24.3) |
| Ventilatory defects<sup>‡</sup>, n (%) |  |
| Obstructive | 27 (77) |
| DL<sub>CO</sub> limited | 17 (49) |
| Lung cancer characteristics |  |
| Lesion size, cm, mean (SD) | 2.4 (1.5) |
| Clinical stage<sup>‡</sup>, n (%) |  |
| IA | 24 (69) |
| IB | 5 (14) |
| IIA | 0 (0) |
| IIB | 3 (9) |
| IIIA | 3 (9) |
| Histology, n (%) |  |
| Adenocarcinoma | 22 (63) |
| Squamous cell carcinoma | 5 (14) |
| Presumed | 8 (23) |
| Primary Treatment, n (%) |  |
| Surgical resection | 18 (51) |
| Definitive radio-ablation | 12 (34) |
| Chemoradiation | 5 (14) |

\*Data available in 31 participants.
<sup>†</sup>Defined as clinical documentation of systolic heart failure or ventricular ejection fraction < 55%.
<sup>‡</sup>Defined as FEV<sub>1</sub>/FVC < 70% for obstructive defect and DL<sub>CO</sub> % predicted < 80 for limitation.
<sup>#</sup>Defined by the 7<sup>th</sup> edition of the American Joint Committee on Cancer TNM staging system.

BMI = body-mass index; CAD = coronary artery disease; CKD = chronic kidney disease; COPD = chronic obstructive pulmonary disease; DL_CO_ = diffusion capacity of the lung for carbon monoxide; FEV1 = forced expiratory volume in 1 second; FVC = forced vital capacity; HFrEF = heart failure with reduced ejection fraction; OSA = obstructive sleep apnea; PTSD = post-traumatic stress disorder; PVD = peripheral vascular disease; SD = standard deviation; TLC = total lung capacity; VASDHS = VA San Diego Healthcare System

### Baseline Functional EC and PRO Assessments

Participants’ baseline functional EC was low [mean 6MWD = 370 m (69% predicted) and impaired in 24 participants (69%) as defined by the lower-limit of normal for healthy adults [31]] (Table 2A). Cancer-specific QoL was also reduced (mean = 72.0 points on scale range 0-100). Approximately half of participants reported abnormal physical function, pain, insomnia, appetite loss, or dyspnea on the EORTC-QLQ-C30/LC-13 (Table 2B) questionnaire (defined as raw scores <mean reference value for functional scales and raw scores >mean reference value for symptom scales [32]). Baseline exploratory outcome assessments are summarized in Table 2C.

**Table 2:**
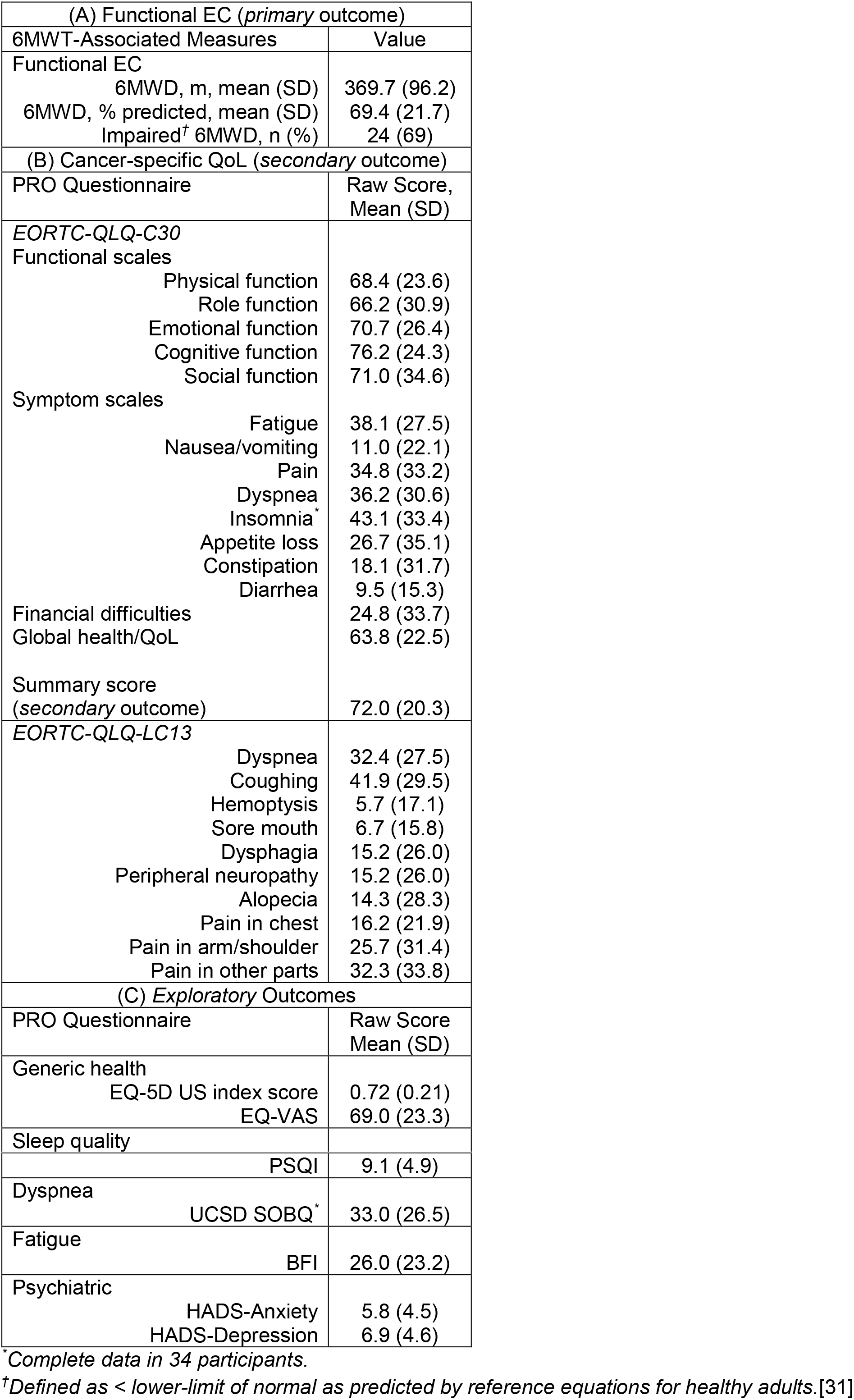
Baseline Functional EC and PRO Assessments (N=35)

| (A) Functional EC (*primary outcome*) |  |
| --- | --- |
| 6MWT-Associated Measures | Value |
| Functional EC |  |
| 6MWD, m, mean (SD) | 369.7 (96.2) |
| 6MWD, % predicted, mean (SD) | 69.4 (21.7) |
| Impaired<sup>†</sup> 6MWD, n (%) | 24 (69) |
| (B) Cancer-specific QoL (*secondary outcome*) |  |
| PRO Questionnaire | Raw Score,<br>Mean (SD) |
| *EORTC-QLQ-C30* |  |
| Functional scales |  |
| Physical function | 68.4 (23.6) |
| Role function | 66.2 (30.9) |
| Emotional function | 70.7 (26.4) |
| Cognitive function | 76.2 (24.3) |
| Social function | 71.0 (34.6) |
| Symptom scales |  |
| Fatigue | 38.1 (27.5) |
| Nausea/vomiting | 11.0 (22.1) |
| Pain | 34.8 (33.2) |
| Dyspnea | 36.2 (30.6) |
| Insomnia<sup>*</sup> | 43.1 (33.4) |
| Appetite loss | 26.7 (35.1) |
| Constipation | 18.1 (31.7) |
| Diarrhea | 9.5 (15.3) |
| Financial difficulties | 24.8 (33.7) |
| Global health/QoL | 63.8 (22.5) |
| Summary score<br>(*secondary outcome*) | 72.0 (20.3) |
| *EORTC-QLQ-LC13* |  |
| Dyspnea | 32.4 (27.5) |
| Coughing | 41.9 (29.5) |
| Hemoptysis | 5.7 (17.1) |
| Sore mouth | 6.7 (15.8) |
| Dysphagia | 15.2 (26.0) |
| Peripheral neuropathy | 15.2 (26.0) |
| Alopecia | 14.3 (28.3) |
| Pain in chest | 16.2 (21.9) |
| Pain in arm/shoulder | 25.7 (31.4) |
| Pain in other parts | 32.3 (33.8) |
| (C) *Exploratory Outcomes* |  |
| PRO Questionnaire | Raw Score<br>Mean (SD) |
| Generic health |  |
| EQ-5D US index score | 0.72 (0.21) |
| EQ-VAS | 69.0 (23.3) |
| Sleep quality |  |

|  |  |  |
| --- | --- | --- |
|  | PSQI | 9.1 (4.9) |
| Dyspnea | UCSD SOBQ * | 33.0 (26.5) |
| Fatigue | BFI | 26.0 (23.2) |
| Psychiatric | HADS-Anxiety | 5.8 (4.5) |
|  | HADS-Depression | 6.9 (4.6) |

\*Complete data in 34 participants.
†Defined as < lower-limit of normal as predicted by reference equations for healthy adults.[31]

6MWD = six-minute walk distance; 6MWT = six-minute walk test; BFI = Brief Fatigue Inventory; EC = exercise capacity; EORTC-QLQ-C30/LC13 = European Organization for Research and Treatment of Cancer QoL Questionnaire Core 30/Lung Cancer Module 13; EQ-5D/VAS = EuroQoL-5 Dimensions/Visual Analogue Scale; HADS = Hospital Anxiety and Depression Scale; PRO = patient-reported outcome; PSQI = Pittsburg Sleep Quality Index; QoL= quality of life; SD = standard deviation; UCSD = University of California San Diego Shortness of Breath Questionnaire; US = United States

### Completion of Follow-Up Assessments

We provided detailed descriptions of curative-intent treatment and treatment-associated morbidities in our online data supplements. Following treatment, 28 (80%) of the 35 participants completed the follow-up 6MWT and 31 (89%) the PRO questionnaires. Two participants had transportation challenges and declined the follow-up 6MWT but completed PRO questionnaires remotely (one via mail and another via telephone). Three participants did not have regular follow-up clinic visits and/or transportation challenges and, therefore, did not complete either 6MWT or PRO re-assessments within the 1-3-month post-treatment period. One participant suffered medical complications following treatment and died during the follow-up period.

### Effect of Treatment on Outcomes

Following curative-intent therapy, there was a statistically non-significant but possibly clinically meaningful decrease in the *primary* outcome functional EC [mean change (95% CI) 6MWD = −25.5 m (−58.4, +7.3), p = 0.12)] (Figure 2A-i), as well nonsignificant change in the *secondary* outcome cancer-specific QoL [EORTC-QLQ-C30 summary score = −3.89 (−10.9, +3.08), p = 0.26)] (Figure 2A-ii) as assessed by the paired-sample t-tests. *Exploratory* outcome assessments showed that dyspnea [mean change (95% CI) UCSD SOBQ = +12.9 (+4.77, +21.0), p = 0.003] (Figure 2A-iii) and fatigue [BFI = +10.4 points (+2.87, +17.9), p = 0.008] (Figure 2A-iv) scores were significantly higher/worse following treatment and no significant change in other *exploratory* outcomes listed in Table 2C.

**Figure 2A:**
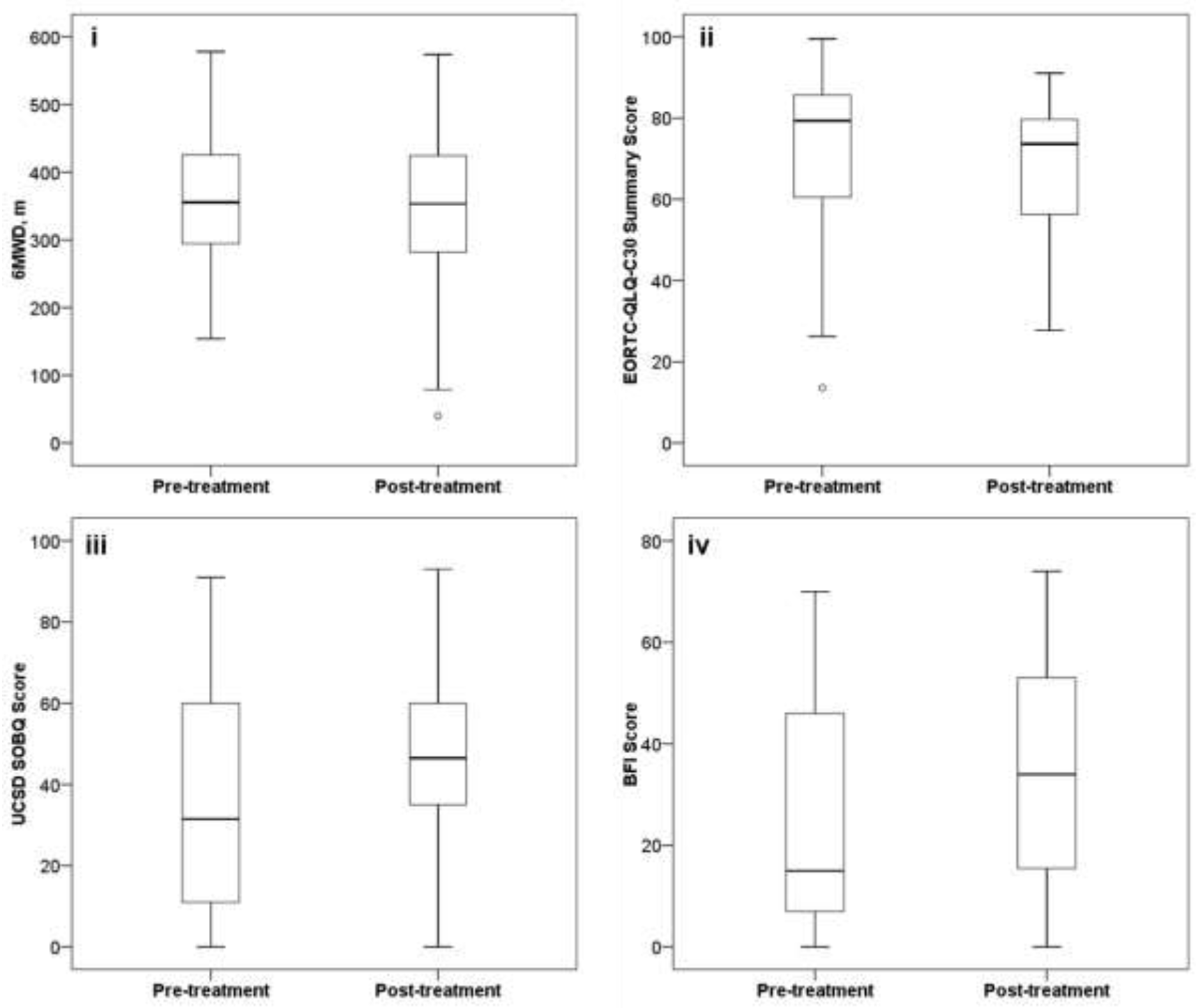
Changes in *Primary, Secondary*, and Significant *Exploratory* Outcomes Associated with Curative-Intent Lung Cancer Treatment. i. Functional EC (*primary* outcome); complete follow-up data in 28 participants; mean difference (post-/pre-treatment) = −25.5 m (95% CI −58.4, +7.29), p = 0.12.
ii. Cancer-specific QoL (*secondary* outcome); complete follow-up data in 31 participants; mean difference (post-/pre-treatment) = −3.89 points (95% CI −10.9, +3.08), p = 0.26.
iii. Dyspnea (UCSD SOBQ, significant *exploratory* outcome) complete data in 30 participants; mean difference (post-/pre-treatment) = +12.9 points (95% CI +4.77, +21.0), p = 0.003.
iv. Fatigue (BFI, significant exploratory outcome); complete data in 31 participants; mean difference (post-/pre-treatment) = +10.4 points (95% CI +2.87, +17.9), p = 0.008.

Results of UVAs and MVAs to identify baseline clinical characteristics associated with the endpoints are shown in E-Tables 2-5. In multivariable GEEs adjusting for all confounders associated with the outcomes, the effect of time (pre-/post-treatment) was borderline statistically associated with a mean 25.4-m decrease in the 6MWD (p = 0.096) (Table 3A) but not cancer-specific QoL (mean summary score 3.39 points decrease, p = 0.30) (Table 3B); dyspnea (mean UCSD SOBQ increase 13.1 points, p = 0.001) (Table 3C) and fatigue (mean BFI increase 9.97 points, p = 0.006) (Table 3D) symptoms were significantly higher/worse following treatment.

**Table 3:** Multivariable GEE Analyses of the Effects of Curative-Intent Lung Cancer Treatment on Outcomes (N=35)

(A) Functional EC (*primary outcome, 6MWD*)
| Variable | $\beta$ (95% CI) | P-value |
| --- | --- | --- |
| Sex (F/M) | 143.5 (101.1, 185.9) | <0.001 |
| Pack year, each | 1.15 (-0.10, 2.39) | 0.07 |
| Hyperlipidemia (N/Y) | 36.82 (-5.39, 79.0) | 0.09 |
| HFrEF (N/Y) | 118.7 (59.6, 177.9) | <0.001 |
| FEV<sub>1</sub>, % predicted | 3.42 (1.70, 5.13) | <0.001 |
| Pack year×FEV<sub>1</sub> % predicted | -0.02 (-0.04, -0.01) | 0.01 |
| **Time effect**<br>**(post-/pre-treatment)** | **-25.4 (-55.3, +4.47)** | **0.096** |

No significant interaction between hyperlipidemia and HFrEF ( $p = 0.98$ ), or pack year and HFrEF ( $p = 0.46$ ). (B) Cancer-specific QoL (*secondary outcome, EORTC-QLQ-C30 summary score*)
| Variable | $\beta$ (95% CI) | P-value |
| --- | --- | --- |
| Smoking status | N/A (F-statistics) | <0.001 |
| HFrEF (N/Y) | 33.9 (23.8, 44.0) | <0.001 |
| Smoking status×HFrEF | N/A | <0.001 |
| FEV<sub>1</sub>, % predicted | 0.18 (-0.03, 0.38) | 0.09 |
| **Time effect**<br>**(post-/pre-treatment)** | **-3.39 (-9.80, +3.02)** | **0.30** |

No significant interaction between smoking status and FEV<sub>1</sub> % predicted ( $p = 0.38$ ). (C) Dyspnea (*significant exploratory outcome, UCSD SOBQ*)
| Variable | $\beta$ (95% CI) | P-value |
| --- | --- | --- |
| Anxiety/depression/PTSD (N/Y) | -22.5 (-33.4, -11.7) | <0.001 |
| FEV<sub>1</sub>, % predicted | -0.39 (-0.62, -0.16) | 0.001 |
| **Time effect**<br>**(post-/pre-treatment)** | **+13.1 (+5.68, +20.6)** | **0.001** |

No significant interaction between anxiety/depression/PTSD and FEV<sub>1</sub> % predicted ( $p = 0.27$ ). (D) Fatigue (*significant exploratory outcome, BFI*)
| Variable | $\beta$ (95% CI) | P-value |
| --- | --- | --- |
| Smoking history | N/A (F-statistics) | <0.001 |
| HFrEF (N/Y) | -15.3 (-30.8, 0.27) | 0.054 |
| Anxiety/depression/PTSD (N/Y) | -36.4 (-56.8, -16.0) | 0.001 |
| Smoking history×<br>Anxiety/depression/PTSD | N/A | 0.001 |
| **Time effect**<br>**(post-/pre-treatment)** | **+9.97 (+2.89, +17.0)** | **0.006** |

For tables A-D: **Bolded** variables indicate time effect (post-/pre-treatment) on outcomes.

6MWD = six-minute walk distance; β = regression coefficient; BFI = Brief Fatigue Inventory; CI = confidence interval; EC = exercise capacity; E0RTC-QLQ-C30 = European Organization for Research and Treatment of Cancer QoL Questionnaire Core 30; F = female; FEV1 = forced expiratory volume in 1 second; GEE = generalized estimating equations; HFrEF = heart failure with reduced ejection fraction; M = male; PTSD = post-traumatic stress disorder; QoL = quality of life; UCSD SOBQ = University of California San Diego Shortness of Breath Questionnaire

In a pre-specified subgroup analysis of stage I patients (N=29) to compare the effects of surgical resection (n=17, 59%) vs definitive radio-ablation (n=12, 41%) on outcomes (Figure 2B), surgical treatment resulted in a significantly higher decrement in the *secondary* outcome cancer-specific QoL following treatment (mean 15.1-point decrease, 95% CI 0.83, 29.4, p =0.04) (Table 4B), but no significant differences in the *primary* outcome functional EC (Table 4A), or *exploratory* outcomes dyspnea or fatigue as assessed by the UCSD SOBQ (Table 4C) or BFI (Table 4D), respectively.

**Table 4:** Subgroup Multivariable GEE Analyses on the Effects of Surgical Treatment Compared to Definitive Radio-ablation on Outcomes in Stage I Patients (N=29)

(A) Functional EC (*primary* outcome, 6MWD)
| Variable | $\beta$ (95% CI) | P-value |
| --- | --- | --- |
| Sex (F/M) | 176.4 (134.1, 218.7) | <0.001 |
| HFrEF (N/Y) | 150.2 (92.6, 207.8) | <0.001 |
| FEV<sub>1</sub> % predicted | 2.97 (1.47, 4.46) | <0.001 |
| Surgical treatment (Y/N) | -41.0 (-111.9, 29.8) | 0.27 |
| Time effect (post-/pre-treatment) | -32.3 (-67.0, 2.46) | 0.07 |
| Surgical treatment × time effect | 9.91 (-56.2, 76.0) | 0.77 |

(B) Cancer-specific QoL (*secondary* outcome, EORTC-QLQ-C30 summary score)
| Variable | $\beta$ (95% CI) | P-value |
| --- | --- | --- |
| Anxiety/depression/PTSD (N/Y) | 15.0 (3.95, 26.0) | 0.01 |
| FEV<sub>1</sub> % predicted | 0.21 (-0.06, 0.47) | 0.12 |
| Surgical treatment (Y/N) | 6.94 (-7.02, 20.9) | 0.33 |
| Time effect (post-/pre-treatment) | 1.34 (-9.82, 12.5) | 0.81 |
| **Surgical treatment × time effect** | **-15.1 (-29.4, -0.83)** | **0.04** |

| Variable | $\beta$ (95% CI) | P-value |
| --- | --- | --- |
| Anxiety/depression/PTSD (N/Y) | -25.3 (-36.6, -14.1) | <0.001 |
| FEV<sub>1</sub> % predicted | -0.24 (-0.52, 0.04) | 0.09 |
| Surgical treatment (Y/N) | -11.8 (-25.6, 2.04) | 0.095 |
| Time effect (post-/pre-treatment) | +16.8 (+9.25, +24.3) | <0.001 |
| Surgical treatment × time effect | 2.37 (-13.2, 17.9) | 0.77 |

(D) Fatigue (*exploratory* outcome, BFI)
| Variable | $\beta$ (95% CI) | P-value |
| --- | --- | --- |
| HFrEF (N/Y) | -21.3 (-39.5, -3.16) | 0.02 |
| Anxiety/depression/PTSD (N/Y) | -24.4 (-37.0, -11.8) | <0.001 |
| FEV<sub>1</sub> % predicted | -0.31 (-0.60, -0.02) | 0.03 |
| Surgical treatment (Y/N) | 6.95 (-6.33, 20.2) | 0.31 |
| Time effect (post-/pre-treatment) | +11.7 (+3.75, +19.6) | 0.004 |
| Surgical treatment × time effect | 5.99 (-9.54, 21.5) | 0.45 |

6MWD = six-minute walk distance; *β* = regression coefficient; BFI = Brief Fatigue Inventory; CI = confidence interval; EC = exercise capacity; EORTC-QLQ-C30 = European Organization for Research and Treatment of Cancer QoL Questionnaire Core 30; F = female; FEV_1_ = forced expiratory volume in 1 second; GEE = generalized estimating equations; M = male; PTSD = post-traumatic stress disorder; QoL = quality of life; UCSD SOBQ = University of California San Diego Shortness of Breath Questionnaire

**Figure 2B:**
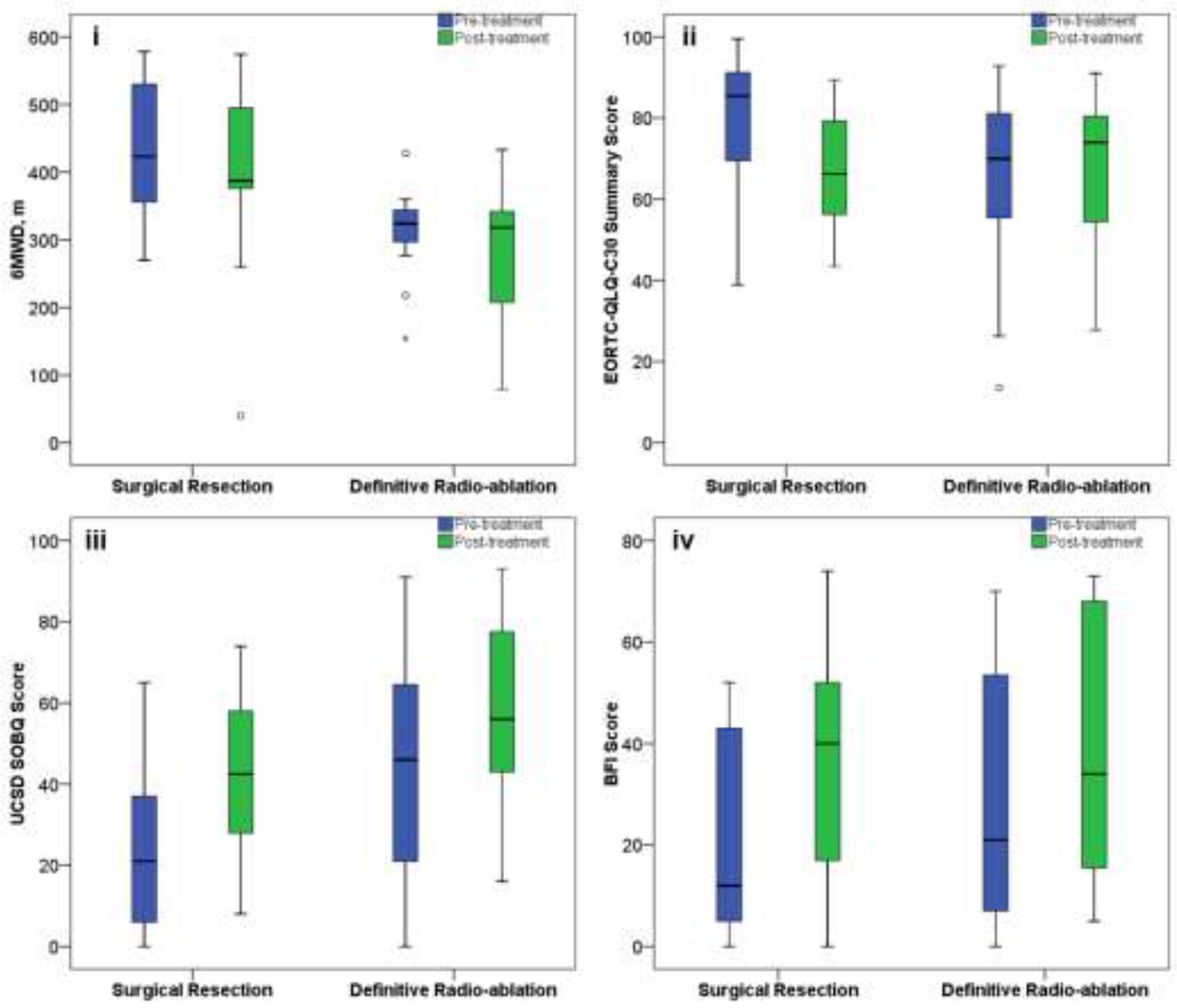
Changes in *Primary, Secondary*, and Significant *Exploratory* Outcomes for Stage I Lung Cancer Stratified by Treatment. i. Functional EC (*primary* outcome); complete response and follow-up in 13 participants for surgical resection and 11 for definitive radio-ablation; no significant between-treatment effect (p = 0.77).
ii. Cancer-specific QoL (*secondary* outcome); complete response and follow-up in 14 participants for surgical resection and 12 for definitive radio-ablation; significant between-treatment effect (p = 0.04).
iii. Dyspnea (UCSD SOBQ, *exploratory* outcome); complete response and follow-up in 14 participants for surgical resection and 12 for definitive radio-ablation; no significant between-treatment effect (p = 0.77). Fatigue (BFI, *exploratory* outcome); complete response and follow-up in 14 participants for surgical resection and 12 for definitive radio-ablation; no significant between-treatment effect (p = 0.45). BFI = brief fatigue inventory; CI = confidence interval; EC = exercise capacity; QoL = quality of life; UCSD SOBQ = University of California San Diego Shortness of Breath Questionnaire

## Discussion

In a prospective, observational cohort study of stage I-IIIA lung cancer patients undergoing curative-intent therapy, we observed: 1) a borderline statistically significant decline in functional EC, the primary outcome; 2) no significant change in the secondary outcome, cancer-specific QoL; and 3) clinically and statistically significant worsening of exploratory outcomes dyspnea and fatigue symptoms at 1 to 3 months following treatment completion.

Much of the literature on the effect of curative-intent lung cancer therapy on health focuses on physiological (i.e., lung function and maximal/peak EC) and clinical (i.e., perioperative morbidity and mortality) outcomes [35, 36]. As such, there is a lack of emphasis on patient-centered outcomes (i.e., functional EC and PROs including QoL) associated with curative-intent therapy [37], which may be more important than survival for some patients [38, 39]. While our study did not detect a statistically significant change in functional EC, the 25-m reduction in the 6MWD following curative-intent therapy is likely clinically significant as suggested by a previous analysis reporting 6MWD changes of 22-42 m as the MCID in the lung cancer population [20].

In addition, to the best of our knowledge, our study is among the first to use the UCSD SOBQ and BFI questionnaires to assess dyspnea and fatigue symptoms, respectively, in patients undergoing curative-intent therapy. Along with the Functional Assessment of Cancer Therapy–General [40]/Lung [41] (FACT-G/L), the EORTC-QLQ-C30 [23]/LC13 [24] questionnaires are the most commonly used cancer/lung cancer-specific PRO instruments. Despite the importance and frequency of abnormal symptoms of dyspnea and fatigue in lung cancer patients [42–45], these two symptoms are assessed by only one and three questions, respectively, in the 30-question EORTC-QLQ-C30, and only three and zero questions, respectively, in the 13-question EORTC-QLQ-LC13. While there are ongoing efforts to create a novel EORTC-QLQ-LC29 instrument with a summary score [46] to more accurately capture lung cancer-specific health, we used separate PRO questionnaires with more questions specifically on these two important symptoms (24 items in the UCSD SOBQ and 9 in the BFI). In this relatively small sample, we detected statistically and clinically meaningful increases/worsening in dyspnea and fatigue symptoms following treatment (as interpreted by their respective MCIDs: 5 points for the UCSD SOBQ [47] and 7 for the BFI [28]).

While lacking long-term follow-up, our study adds to existing literature on the effects of curative-intent lung cancer treatment on PROs and QoL. Among the largest studies to date including 156 consecutive lung cancer patients, Brunelli and coworkers [48] reported that in those undergoing major lung resection surgery, the physical composite scale in the generic QoL Short Form-36 Health Survey was significantly reduced at one month but completely recovered at three months, and the mental composite scale remained unchanged. In the same study, the authors also found poor correlation (coefficients <0.2) between these generic health measures and FEV1, DLco, and EC as assessed by the height reached on the stair-climbing test, and suggested that functional variables cannot substitute for QoL measures [48]. In contrast, in a smaller study but with longer two-year follow-up, Ilonen and coworkers [49] observed that in 53 patients undergoing major lung cancer resection surgery, the generic QoL as assessed by the 15-Dimensions index score was decreased compared to preoperative values at 3, 12, and 24 months following surgery. They also found no correlation between preoperative FEV_1_ or DL_CO_ and QoL at any of the follow-up assessment time points [49]. Similarly, in a prospective cohort study of 131 lung cancer patients undergoing lobectomy or bilobectomy, Schulte and coworkers [50] found that most domains of health as assessed by the E0RTC-QLQ-C30/LC13, including physical function, pain, and dyspnea, were significantly impaired after surgery and remained so for up to 24 months following treatment. Of note, each health domain (9 psychometric and 28 single-item scales) was analyzed separately in this study [50].

While these contrasting findings may partly be due to differences in baseline lung function and/or clinical lung cancer stage which are not completely reported in all three studies, standard physiological outcome assessments including pulmonary function testing do not appear to adequately capture all the effects of curative-intent lung cancer treatment on health including PROs and QoL. Also, these studies suggest that results may vary depending on the PRO or QoL instruments used, possibly due to a lack of a validated questionnaire for lung cancer patients undergoing curative-intent therapy, variations in psychometric properties between instruments including sensitivity to change, or the availability of a composite score to avoid multiple testing and minimize chance bias.

A recent exploratory analysis of a RCT involving 22 stage IA NSCLC patients to investigate the effects of surgical resection compared to SBRT on PROs and QoL suggested that SBRT may be more favorable than lobectomy, specifically in the EORTC-QLQ-C30/LC13 subscale global health status/QoL using univariable analysis of time to clinically significant deterioration (hazard ratio = 0.19, 95% CI 0.04-0.91, p = 0.038) [51]. To the best of our knowledge, our subgroup analysis is among the first to use the validated composite EORTC-QLQ-C30 summary score [22] in stage I lung cancer patients to prospectively investigate the effects of surgical vs definitive radio-ablative therapy on cancer-specific QoL. Our multivariable analysis suggests that surgical resection may lead to greater decrement in cancer-specific QoL compared to definitive radio-ablative therapy at 1 to 3 months following treatment. While the sample sizes are small and selected, these findings have implications in the shared-decision making, treatment selection, and/or post-treatment care in this subgroup of patients.

While adult cancer survivors experience both physical and psychological impairments, more may have reduced QoL as the result of physical than psychological impairments which can go undetected and untreated and result in disability [52]. Compared to usual care, physical exercise has been shown to be effective in improving QoL and physical function in cancer survivors based on a recent systematic review of 34 randomized clinical trials (RCTs) [53]. However, out of the 34 RCTs examined, only one small study (N=51) focused on lung cancer survivors [53]. While highlighting the importance of exercise to improve QoL and function in cancer survivors, these findings also suggest caution in efforts to improve lung cancer survivorship. Among them, lung cancer patients are somewhat different from those with other cancers due to higher median age at diagnosis, lifetime exposure to tobacco, and prevalence of pulmonary and cardiovascular comorbidities [15]. In addition, the effects of curative-intent lung cancer treatment have unique effects related to cardiopulmonary health since part of the therapy requires local destruction and/or removal of lung tissue which may impede cardiopulmonary function and EC in some. Our exploratory PRO assessments suggest that decreasing symptom burden due to dyspnea and fatigue (e.g. through optimizing medical therapy for cardiopulmonary disease) may be important to improve exercise, function, and/or QoL in these patients.

Our study has limitations. First, the relatively small sample size may not be adequately powered to detect statistically significant differences in the primary or secondary outcomes. Second, the range of 1 to 3 months for follow-up assessments may lead to additional variations in the outcomes measured and, therefore, diminished statistical power. Third, the absence of long-term follow-up assessments limits our ability to draw conclusions on the effect of time on outcomes including exploratory variables following treatment. For instance, it is possible that some of the worsening in dyspnea and fatigue may improve spontaneously after the 3-month follow-up period. Finally, our findings may have limited generalizability due to it being a single-institutional study involving a predominantly white male veteran patient population with a significant tobacco exposure and higher than expected prevalence of comorbidities including coronary artery disease and COPD [15].

The strengths of our study include pre-specified primary, secondary, and exploratory outcomes to minimize chance bias. We collected a thorough list of potential confounders, all of which were entered in the electronic medical record and collected by board-certified physicians, maximizing the accuracy of the data obtained. In addition, all baseline and most follow-up functional EC and PRO assessments were performed in-person by one observer (DH), maximizing the completeness and accuracy of the data collected and minimizing inter-observer variability. Equally important, we had a high completion rate (at least 80%) on all outcomes measured, maximizing the validity of our findings. In addition, unlike many of the published studies to date, we used multivariable GEE analyses to adjust for baseline characteristics including lung function associated with the outcomes, enhancing our conclusions. Finally, we provided detailed descriptions of important clinical events following curative-intent lung cancer treatment and interpreted outcome changes using MCIDs where available, facilitating translation to the clinical setting.

We conclude that in a hypothesis-generating, prospective observational cohort study of lung cancer patients undergoing curative-intent therapy, there were statistically significant and clinically meaningful worsening of dyspnea and fatigue symptoms, possible decreases in functional EC, but no significant change in cancer-specific QoL at 1 to 3 months following treatment. In stage I lung cancer patients, surgical treatment may lead to a greater decrement in cancer-specific QoL compared to definitive radio-ablative therapy. These results provide a proof-of-concept on the information provided by physio-psychological assessments in this patient population and may facilitate future studies to reduce symptom burden, and/or improve functional EC and QoL.

## Acknowledgements

We would like to thank Dr. Florin Vaida and Anya Umlauf for assistance in the statistical analyses for this study, the Division of Pulmonary, Critical Care, and Sleep Medicine and the Clinical and Translational Research Institute (CTRI) Clinical Research Enhancement through Supplemental Training (CREST)/Master of Advanced Study (MAS) in Clinical Research Program at the University of California San Diego (UCSD) for financial and scholarship support, the Section of Pulmonary and Critical Care Medicine at the VA San Diego Healthcare System (VASDHS), notably Dr. Judd Landsberg and Dr. Philippe Montgrain for their relentless support, Svetlana Sheinkman for maintaining an ongoing list of lung cancer patients, Christine Miller for assistance with patient recruitment and enrollment, and the staff in the Pulmonary Function Laboratory for assistance with patient scheduling and six-minute walk test assessments. We would also like to give tremendous thanks to all the veteran lung cancer survivors who volunteered and donated their time for this study.

## Abbreviations

6MWD: six-minute walk distance
6MWT: six-minute walk test
AECOPD: acute exacerbation of COPD
ATS: American Thoracic Society
BFI: Brief Fatigue Inventory
CI: confidence interval
COPD: chronic obstructive pulmonary disease
CTB: chest tumor board
DL_CO_: diffusion capacity of the lung for carbon monoxide
EC: exercise capacity
EORTC-QLQ-C30/LC13: European Organization for Research and Treatment of Cancer QoL Questionnaire Core 30/Lung Cancer Module 13
EQ-5D/VAS: EuroQoL-5 Dimensions/visual analogue scale
FACT-G/L: Functional Assessment of Cancer Therapy – General/Lung
FEV1: forced expiratory volume in 1 second
GEE: generalized estimating equations
HADS: Hospital Anxiety and Depression Scale
HF: heart failure
LCS: lung cancer screening
MCID: minimal clinically important difference
MVA: multivariable linear regression analysis
NSCLC: non-small cell lung cancer
PRO: patient-reported outcome
PSQI: Pittsburgh Sleep Quality Index
QoL: quality of life
RCT: randomized clinical trial
RFA: radiofrequency ablation
SBRT: stereotactic body radiotherapy
TLC: total lung capacity
UCSD SOBQ: University California San Diego Shortness of Breath Questionnaire
US: United States
UVA: univariable linear regression analysis
VASDHS: VA San Diego Healthcare System

## Notes

Funding: This work was supported by the National Institutes of Health (L30CA208950 and 1T32HL134632-01); and the American Cancer Society (PF-17-020-01-CPPB).

Conflict of Interest: All authors declare that there is no conflict of interest.

