## Supplemental Data for "Effects of Curative-Intent Lung Cancer Therapy on Functional Exercise Capacity and Patient-Reported Outcomes"

**ONLINE DATA SUPPLEMENTS**

**Methods**

*Confounders:* We collected baseline clinical characteristics and potential confounders important in lung cancer and cardiopulmonary health and QoL, including age, sex, body-mass index, tobacco exposure, comorbidities (including COPD, HF, and psychiatric illness), lung function [FEV_1_, total lung capacity (TLC), DL_CO_], and echocardiographic findings where available; lung cancer-related information included histologic subtype, clinical stage, and primary treatment modality.

**Results**

*Curative-Intent Treatment:* All but two participants underwent either surgical resection, definitive radio-ablative therapy, or concurrent chemoradiation for treatment. Of the 18 (51%) participants who underwent surgical resection, all but two received lobectomy; one underwent pneumonectomy due a central tumor location, and another wedge resection due to poor lung function and planned stereotactic body radiotherapy (SBRT) for a synchronous primary lung cancer (follow-up outcome assessments were obtained after wedge resection in this participant); no participant received adjuvant chemo- or radiotherapy. Among the 12 (34%) participants who underwent definitive radio-ablative therapy, all but two received SBRT; one received cryoablation due to a history of pneumonitis following radiotherapy for a previous primary lung cancer, and another received radiofrequency ablation (RFA) and SBRT for synchronous primary lung cancers (follow-up outcome assessments were obtained following completion of SBRT and RFA). Of the 5 (14%) participants undergoing concurrent chemoradiation, two received tri-modality therapy (follow-up assessments were performed at 1 to 3 months following completion lobectomy in these participants).

*Treatment-Associated Morbidities*: Following lobectomy, one participant had recurrent mixed hypoxemic/hypercarbic respiratory failure twice requiring intubation with mechanical ventilation and a 29-day hospital stay with clinical documentation of volume overload, lobar collapse, pneumonia, persistent air leak requiring chest tube re-insertion, deep vein thrombosis, and acute kidney injury in the postoperative period. Another participant had a 10-day hospital stay related to subcutaneous emphysema and persistent air leak requiring chest tube insertion. A third participant had a 5-day hospital stay due to concern of intrathoracic bleeding and was discharged with a walker to assist with activities of daily living (follow-up 6MWT was performed with the walker; no walker was needed in the pre-treatment assessment). Following definitive radio-ablative therapy, one participant had radiation pneumonitis requiring oral corticosteroids and supplemental oxygen therapy. Another had pneumothorax following RFA requiring chest tube insertion and hospitalization and subsequent re-admission for dyspnea requiring oral corticosteroids and antibiotics for acute exacerbation of COPD (AECOPD) (follow-up assessments were performed after these events). Another participant had AECOPD requiring medical treatment and hospitalization at approximately 2 months following SBRT (follow-up assessments were performed at approximately 3 weeks after hospital discharge). Following concurrent chemoradiation, one participant died with documented diagnoses of neutropenia, pneumonia, sepsis, and pneumonitis; another was too debilitated to perform follow-up 6MWT but able to complete PRO assessments at approximately 3 months following treatment.

E-Table 1: Comparison of Clinical Characteristics of Participants (N=35) vs. Non-Participants (N=15)

| *Clinical Characteristic* | *Difference*  *Mean (SE)* | *P-value^†^* |
| --- | --- | --- |
| Age, year | -3.40 (2.32) | 0.15 |
| Sex | N/A | 1.00 |
| White race | N/A | 0.49 |
| BMI, kg/m^2^ | -0.23 (1.56) | 0.88 |
| Pack years | 0.95 (9.88) | 0.92 |
| Comorbidities  Hypertension  Hyperlipidemia  Diabetes  CKD  Atrial fibrillation/flutter  CAD  HFrEF  PVD  COPD  OSA  Anxiety/depression/PTSD  Other cancer | N/A  N/A  N/A  N/A  N/A  N/A  N/A  N/A  N/A  N/A  N/A  N/A | 0.73  0.54  0.25  1.00  0.70  0.75  0.65  0.46  0.30  0.30  0.73  0.75 |
| Lung function*^*^*  FEV_1_/FVC, %  FEV_1_, % predicted  TLC, % predicted  DL_CO_, % predicted | 7.57 (4.87)  9.00 (8.14)  -5.89 (7.40)  14.9 (7.72) | 0.13  0.27  0.43  0.06 |
| Lung cancer characteristics  **Stage I disease**  Surgical treatment | **N/A**  N/A | **0.04**  0.55 |

*^*^Data available in 13, 7, and 12 non-participants for FEV_1_, TLC, and DL_CO_ % predicted, respectively.*

*^†^Independent sample t-tests for continuous variables and Fisher’s exact tests for categorical variables.*

***Bolded*** *variable indicates significant differences in proportions of stage II-IIIA disease: 6/35 (17%) for participants vs 7/15 (47%) for non-participants.*

BMI = body-mass index; CAD = coronary artery disease; CKD = chronic kidney disease; COPD = chronic obstructive pulmonary disease; DL_CO_ = diffusion capacity of the lung for carbon monoxide; FEV_1_ = forced expiratory volume in 1 second; FVC = forced vital capacity; HFrEF = heart failure with reduced ejection fraction; OSA = obstructive sleep apnea; PTSD = post-traumatic stress disorder; PVD = peripheral vascular disease; SE = standard error; TLC = total lung capacity.

E-Table 2A: UVA – Predictors of 6MWD

| *Variable* | *β (95% CI)* |
| --- | --- |
| Age, year | -2.54 (-7.11, 2.03) |
| White race (N/Y) | 31.1 (-48.1, 110.3) |
| BMI, kg/m^2^ | 3.96 (-3.03, 11.0) |
| **Sex (F/M)** | **165.1 (-27.8, 358.0)** |
| Tobacco exposure  Smoking status  **Pack years** | N/A (F-statistics)  **-0.97 (-1.91, -0.04)** |
| Hypertension (N/Y) | 26.8 (-46.9, 100.5) |
| **Hyperlipidemia (N/Y)** | **44.1 (-22.6, 110.9)** |
| Diabetes (N/Y) | 5.22 (-71.6, 82.0) |
| CKD (N/Y) | 13.6 (-82.2, 109.4) |
| Atrial fibrillation/flutter (N/Y) | 23.7 (-55.8, 103.2) |
| **CAD (N/Y)** | **51.2 (-18.8, 121.2)** |
| **HFrEF (N/Y)** | **102.0 (13.2, 190.9)** |
| PVD (N/Y) | 38.6 (-49.4, 126.6) |
| **COPD (N/Y)** | **85.9 (18.1, 153.7)** |
| OSA (N/Y) | -56.0 (-159.6, 47.6) |
| Anxiety/Depression/PTSD (N/Y) | 16.6 (-60.0, 93.2) |
| Other cancer (N/Y) | 11.0 (-63.2, 85.2) |
| Lung function  **FEV_1_, % predicted**  TLC, % predicted  **DL_CO_, % predicted**  Ventilatory defect  Obstructive defect (N/Y)  DL_CO_ limited (N/Y) | **2.12 (0.92, 3.31)**  -0.60 (-2.40, 1.21)  **2.00 (0.79, 3.21)**  78.2 (3.23, 153.2)  82.7 (22.3, 143.2) |
| Stage I (N/Y) | -24.7 (-113.4, 63.9) |
| **Presumed lung cancer (N/Y)** | **75.9 (0.58, 151.2)** |
| **Treatment group** | **N/A (F-statistics)** |

***Bolded*** *variables indicate associations at p < 0.20 and therefore selected to enter MVAs.*

E-Table 2B: MVA – Independent Predictors of 6MWD

| *Variable* | *β (95% CI)* | *P-value* |
| --- | --- | --- |
| Sex (F/M) | 113.8 (-34.3, 261.8) | 0.13 |
| Pack year, each | -0.56 (-1.34, 0.21) | 0.15 |
| Hyperlipidemia (N/Y) | 42.8 (-8.33, 93.9) | 0.10 |
| HFrEF (N/Y) | 90.8 (21.7, 160.0) | 0.01 |
| FEV_1_, % predicted | 1.95 (0.87, 3.02) | 0.001 |

*Overall model R^2^ = 0.56, p < 0.001.*

6MWD = six-minute walk distance; β = regression coefficient; BMI = body-mass index; CAD = coronary artery disease; CI = confidence interval; CKD = chronic kidney disease; COPD = chronic obstructive pulmonary disease; DL_CO_ = diffusion capacity of the lung for carbon monoxide; FEV_1_ = forced expiratory volume in 1 second; FVC = forced vital capacity; HFrEF = heart failure with reduced ejection fraction; MVA = multivariable linear regression analysis; OSA = obstructive sleep apnea; PTSD = post-traumatic stress disorder; PVD = peripheral vascular disease; SE = standard error; TLC = total lung capacity; UVA = univariable linear regression analysis

E-Table 3A: UVA – Predictors of EORTC-QLQ-C30 Summary Score

| *Variable* | *β (95% CI)* |
| --- | --- |
| Age, year | 0.45 (-0.52, 1.42) |
| White race (N/Y) | -2.44 (-19.3, 14.4) |
| **BMI, kg/m^2^** | **1.13 (-0.32, 2.58)** |
| Sex (F/M) | 14.2 (-28.0, 56.5) |
| Tobacco exposure  **Smoking status**  Pack years | **N/A (F-statistics)**  -0.02 (-0.23, 0.19) |
| Hypertension (N/Y) | -3.65 (-19.3, 12.0) |
| Hyperlipidemia (N/Y) | 6.18 (-8.12, 20.5) |
| Diabetes (N/Y) | -3.58 (-19.7, 12.6) |
| CKD (N/Y) | 0.57 (-19.7, 20.8) |
| Atrial fibrillation/flutter (N/Y) | 6.66 (-10.0, 23.4) |
| CAD (N/Y) | -1.10 (-16.4, 14.2) |
| **HFrEF (N/Y)** | **27.8 (10.1, 45.5)** |
| PVD (N/Y) | -7.47 (-26.1, 11.1) |
| COPD (N/Y) | 9.54 (-5.78, 24.9) |
| OSA (N/Y) | -7.76 (-29.9, 14.3) |
| **Anxiety/Depression/PTSD (N/Y)** | **14.1 (-1.31, 29.5)** |
| Other cancer (N/Y) | 3.46 (-12.2, 19.1) |
| Lung function  **FEV_1_, % predicted**  TLC, % predicted  DL_CO_, % predicted  Ventilatory defect  Obstructive defect (N/Y)  DL_CO_ limited (N/Y) | **0.22 (-0.07, 0.51)**  -0.16 (-0.49, 0.17)  0.14 (-0.15, 0.44)  9.61 (-6.92, 26.1)  7.59 (-6.32, 21.5) |
| Stage I (N/Y) | -7.48 (-26.1, 11.1) |
| Presumed lung cancer (N/Y) | -1.99 (-18.8, 14.9) |
| **Treatment group** | **N/A (F-statistics** |

***Bolded*** *variables indicate associations at p < 0.20 and therefore selected to enter MVAs.*

E-Table 3B: MVA – Independent Predictors of EORTC-QLQ-C30 Summary Score

| *Variable* | *β (95% CI)* | *P-value* |
| --- | --- | --- |
| Smoking status | N/A (F-statistics) | 0.18 |
| HFrEF (N/Y) | 30.1 (12.8, 47.4) | 0.001 |
| Anxiety/depression/PTSD (N/Y) | 12.8 (-1.10, 26.7) | 0.07 |
| FEV_1_, % predicted | 0.21 (-0.10, 0.53) | 0.18 |
| Treatment group | N/A (F-statistics) | 0.10 |

*Overall model R^2^ = 0.59, p = 0.001.*

β = regression coefficient; BMI = body-mass index; CAD = coronary artery disease; CI = confidence interval; CKD = chronic kidney disease; COPD = chronic obstructive pulmonary disease; DL_CO_ = diffusion capacity of the lung for carbon monoxide; EORTC-QLQ-C30 = European Organization for Research and Treatment of Cancer QoL Questionnaire Core 30; FEV_1_ = forced expiratory volume in 1 second; FVC = forced vital capacity; HFrEF = heart failure with reduced ejection fraction; MVA = multivariable linear regression analysis; OSA = obstructive sleep apnea; PTSD = post-traumatic stress disorder; PVD = peripheral vascular disease; SE = standard error; TLC = total lung capacity; UVA = univariable linear regression analysis

E-Table 4A: UVA – Predictors of UCSD SOBQ

| *Variable* | *β (95% CI)* |
| --- | --- |
| Age, year | -0.19 (-1.47, 1.10) |
| White race (N/Y) | -5.92 (-28.0, 16.1) |
| BMI, kg/m^2^ | -1.07 (-3.08, 0.93) |
| Sex (F/M) | -33.0 (-87.3, 21.3) |
| Tobacco exposure  **Smoking status**  Pack years | **N/A (F-statistics)**  0.18 (-0.09, 0.45) |
| Hypertension (N/Y) | 10.7 (-9.54, 31.0) |
| Hyperlipidemia (N/Y) | -5.64 (-24.6, 13.4) |
| Diabetes (N/Y) | 6.84 (-14.3, 28.0) |
| CKD (N/Y) | -2.08 (-28.6, 24.4) |
| Atrial fibrillation/flutter (N/Y) | -7.97 (-29.9, 14.0) |
| CAD (N/Y) | 5.42 (-14.6, 25.4) |
| HFrEF (N/Y) | -7.94 (-34.3, 18.4) |
| **PVD (N/Y)** | **16.2 (-7.72, 40.2)** |
| **COPD (N/Y)** | **-21.9 (-40.9, -2.79)** |
| OSA (N/Y) | 15.6 (-13.0, 44.2) |
| **Anxiety/Depression/PTSD (N/Y)** | **-28.2 (-46.9, -9.50** |
| Other cancer (N/Y) | 6.70 (-13.8, 27.2) |
| Lung function  **FEV_1_, % predicted**  **TLC, % predicted**  **DL_CO_, % predicted**  Ventilatory defect  Obstructive defect (N/Y)  DL_CO_ limited (N/Y) | **-0.50 (-0.85, -0.14)**  **0.37 (-0.11, 0.85)**  **-0.32 (-0.71, 0.07)**  -20.5 (-41.4, 0.42)  -13.9 (-32.1, 4.17) |
| Stage I (N/Y) | 8.17 (-18.2, 34.5) |
| Presumed lung cancer (N/Y) | -4.87 (-27.0, 17.2) |
| **Treatment group** | **N/A (F-statistics)** |

***Bolded*** *variables indicate associations at p < 0.20 and therefore selected to enter MVAs.*

E-Table 4B: MVA – Independent Predictors of UCSD SOBQ

| *Variable* | *β (95% CI)* | *P-value* |
| --- | --- | --- |
| PVD (N/Y) | 15.8 (-3.67, 35.3) | 0.11 |
| Anxiety/Depression/PTSD (N/Y) | -30.0 (-46.4, -13.6) | 0.001 |
| Treatment group | N/A (F-statistics) | 0.047 |
| FEV_1_, % predicted | -0.36 (-0.74, 0.02) | 0.06 |

*Overall model R^2^ = 0.55, p < 0.001.*

β = regression coefficient; BMI = body-mass index; CAD = coronary artery disease; CI = confidence interval; CKD = chronic kidney disease; COPD = chronic obstructive pulmonary disease; DL_CO_ = diffusion capacity of the lung for carbon monoxide; FEV_1_ = forced expiratory volume in 1 second; FVC = forced vital capacity; HFrEF = heart failure with reduced ejection fraction; MVA = multivariable linear regression analysis; OSA = obstructive sleep apnea; PTSD = post-traumatic stress disorder; PVD = peripheral vascular disease; SE = standard error; TLC = total lung capacity; UCSD SOBQ = University of California San Diego Shortness of Breath Questionnaire; UVA = univariable linear regression analysis

E-Table 5: UVA – Predictors of BFI

| *Variable* | *β (95% CI)* |
| --- | --- |
| **Age, year** | **-0.74 (-1.83, 0.36)** |
| White race (N/Y) | 3.37 (-15.9, 22.7) |
| BMI, kg/m^2^ | -0.99 (-2.68, 0.69) |
| Sex (F/M) | -21.6 (-69.7, 26.5) |
| Tobacco exposure  **Smoking status**  Pack years | **N/A (F-statistics)**  0.10 (-0.14, 0.34) |
| **Hypertension (N/Y)** | **11.9 (-5.61, 29.3)** |
| Hyperlipidemia (N/Y) | -5.17 (-21.6, 11.3) |
| Diabetes (N/Y) | 9.31 (-8.96, 27.6) |
| CKD (N/Y) | 7.50 (-15.5, 30.5) |
| Atrial fibrillation/flutter (N/Y) | -4.01 (-23.3, 15.3) |
| CAD (N/Y) | 7.20 (-10.1, 24.5) |
| **HFrEF (N/Y)** | **-14.7 (-37.3, 7.93)** |
| **PVD (N/Y)** | **17.9 (-2.64, 38.5)** |
| COPD (N/Y) | -11.1 (-28.6, 6.43) |
| OSA (N/Y) | 15.8 (-9.04, 40.7) |
| **Anxiety/Depression/PTSD (N/Y)** | **-26.4 (-42.5, -10.4)** |
| Other cancer (N/Y) | 3.54 (-14.4, 21.5) |
| Lung function  FEV_1_, % predicted  TLC, % predicted  DL_CO_, % predicted  Ventilatory defect  Obstructive defect (N/Y)  DL_CO_ limited (N/Y) | -0.20 (-0.53, -0.14)  0.15 (-0.29, 0.59)  -0.13 (-0.46, 0.21)  -6.84 (-26.0, 12.3)  -7.04 (-23.1, 9.01) |
| Stage I (N/Y) | 10.6 (-10.6, 31.8) |
| Presumed lung cancer (N/Y) | 9.11 (-9.94, 28.2) |
| Treatment group | N/A (F-statistics) |

***Bolded*** *variables indicate associations at p < 0.20 and therefore selected to enter MVAs.*

E-Table 5B: MVA – Independent Predictors of BFI

| *Variable* | *β (95% CI)* | *P-value* |
| --- | --- | --- |
| Smoking history | N/A (F-statistics) | 0.03 |
| HFrEF (N/Y) | -20.9 (-39.2, -2.46) | 0.03 |
| PVD (N/Y) | 19.8 (2.02, 37.7) | 0.03 |
| Anxiety/depression/PTSD (N/Y) | -13.2 (-29.6, 3.20) | 0.11 |

*Overall model R^2^ = 0.51, p = 0.001.*

β = regression coefficient; BFI = Brief Fatigue Inventory; BMI = body-mass index; CAD = coronary artery disease; CI = confidence interval; CKD = chronic kidney disease; COPD = chronic obstructive pulmonary disease; DL_CO_ = diffusion capacity of the lung for carbon monoxide; FEV_1_ = forced expiratory volume in 1 second; FVC = forced vital capacity; HFrEF = heart failure with reduced ejection fraction; MVA = multivariable linear regression analysis; OSA = obstructive sleep apnea; PTSD = post-traumatic stress disorder; PVD = peripheral vascular disease; SE = standard error; TLC = total lung capacity; UVA = univariable linear regression analysis
